# Legacy effect of microplastics on plant-soil feedbacks

**DOI:** 10.1101/2022.04.12.488083

**Authors:** Y.M Lozano, M.C Rillig

## Abstract

Microplastics are a complex contaminant suite that are now understood to affect plants and soil biota and the processes they drive. However, the role of microplastic in plant-soil feedbacks, a key feature in plant-soil interactions, is still unknown. We address this here, using soil from a previous experiment, which has been conditioned with 12 different microplastic types including fibers, films, foams, and fragments. To evaluate the feedback effect, we grew a native and a range-expanding plant species with inocula extracted from each one of these soils. At harvest, plant biomass and root morphological traits were measured.

Films gave rise to a positive feedback on shoot mass (higher mass with soil inocula conditioned with microplastics than without), likely via negative effects on harmful soil biota. Foams and fragments also caused positive feedback on shoot mass likely via effects on enzymatic activities and mutualistic soil biota. Fibers led to negative feedback on root mass as they may promote the abundance of soil pathogens.

Microplastics also have a legacy effect on root traits: *Daucus* had thicker roots probably for promoting mycorrhizal associations while *Calamagrostis* had reduced root diameter probably for diminishing pathogenic infection. Microplastic legacy on plants is species-specific and may affect plant biomass primarily via root traits. Microplastics, as a function of their shape and polymer type, have a feedback effect on plant performance.

## Introduction

Microplastics (<5 mm) are an important new global change factor with a high potential to affect terrestrial ecosystems worldwide (Rillig, 2012; De Souza Machado et al., 2018; Lozano et al., 2021a). These particles may appear in different shapes (e.g., fibers, films, foams, fragments) and sizes, spanning a wide range of physical and chemical properties (Helmberger et al., 2020). Microplastics (MPs) can pollute the soil through different pathways such as soil amendments, plastic mulching, irrigation, flooding, atmospheric input, littering, and street runoff (Rillig, 2012; Bläsing and Amelung, 2018; De Souza Machado et al., 2018; Brahney et al., 2020), causing recognized effects on soil properties and soil biota.

Microplastics as a function of their shape may affect soil aggregation, soil bulk density, nutrient retention, or soil pH (De Souza Machado et al., 2019, Lozano et al., 2021ab, Zhao et al., 2021). In the shape of films, MPs may create additional channels for water movement, increasing the rate of soil evaporation (Wan et al., 2019); while by contrast, in the shape of fibers MPs may hold water in the soil for longer, enhancing soil water content (De Souza Machado et al., 2019, Lozano et al., 2021b). In addition, microplastic foams or fragments may increase soil porosity and aeration (Carter and Gregorich, 2006; Ruser et al., 2008). All these microplastic effects on soil properties may in turn have consequences for soil biota. Indeed, MPs as a function of their shape may affect soil microbial respiration, enzymatic activity, or soil microbial community composition (Lozano et al., 2021b; Zhao et al., 202, Fei et al., 2020).

The polymer type from which the microplastic is made also affects soil properties and thus soil biota (Lozano et al., 2021b). In order to prolong plastic life and enhance polymer properties such as flexibility, durability, color, or resistance (Hahladakis et al., 2018; Waldman and Rillig 2020), additives such as light and thermal stabilizers, UV absorbers, colored pigments, anti-fog substances, and antioxidants (Hahladakis et al., 2018) are used in plastic manufacture. Many of these additives may potentially leach into the soil with harmful consequences and toxic effects on soil biota. Negative effects of microplastic have been detected on nematode reproduction (Kim et al., 2020), earthworm performance (Huerta Lwanga et al., 2016), as well as on soil microorganisms. In fact, microplastics may cause a decline in soil bacterial diversity and richness (Huang et al., 2019; Fei et al., 2020), and also potentially cause negative effects on soil fungal communities (Kettner et al., 2017; Leifheit et al., 2021; Zhu et al., 2022).

Previous research shows that microplastics may promote shoot and root mass as a function of their shape and polymer type (Lozano et al., 2021b), responses that have been linked to microplastic effects on soil properties, as for example, the increase of soil macroporosity (Carter and Gregorich, 2006; Ruser et al., 2008), which facilitates root penetration and thus plant growth. By contrast, microplastics could also cause a reduction in plant growth if they, for instance, contain toxic substances (van Kleunen et al., 2019). These positive or negative effects of microplastics on plant performance can also be explained by plant-soil feedback. That is, the microplastic legacy effect on soil biota and their consequences or feedback on plant performance; a phenomenon that may occur if the microplastic is present in the soil or if it is removed (e.g., after being degraded or transported down in the soil profile). Microplastics in soil can feedback plant species either positively, negatively, or neutrally (sensu Bever et al., 1997). Positive feedback (an increase in plant performance driven by microplastic legacy) or negative feedback (the opposite) can be linked with the accumulation of soil biota that can improve plant performance, such as beneficial rhizosphere bacteria, fixing nitrogen bacteria, mutualistic fungi (Bever et al., 1997), or with the accumulation of pathogens that can suppress plant growth, such as pathogenic bacteria and fungi or nematodes (Bever et al., 1997, Lithner et al., 2011, van der Putten et al., 2016, Hahladakis et al., 2018, Bennett and Klironomos, 2019).

Microplastics’ legacy on soil would also create new habitat conditions that may favor some plant species. In that regard, previous research has shown that microplastics may promote the growth of species of range-expanding character over other native species (Lozano and Rillig, 2020), however, the mechanisms underlying this phenomenon are still unclear. One possibility is that the microplastic legacy on soil biota may positively feedback on species of invasive character while negatively on other native species. This, as mechanisms that promote plant invasiveness such as facilitation by soil biota or release of natural enemies (Daneshgar and Jose, 2009) would be potentially promoted by the microplastics legacy on soil biota, which in the end may help explain the competitive success of species of invasive character in novel environments.

The implications of the legacy effect of microplastics in soil have not yet been elucidated, despite interest in legacy effects of global change factors (Meisner et al., 2013, Duell et al., 2019). Thus, we hypothesize that the legacy effect of microplastics on soil would affect the magnitude and direction of the soil feedback on plants depending on the microplastic shape and polymer type with which the soil was conditioned, as well as on the plant trait studied. We included root traits since in addition to plant biomass, root morphological traits are key indicators of plant-soil feedback responses to global change factors (Meisner et al., 2013). Likewise, our study aims to understand whether the legacy effect of microplastics on soil may contribute to the competitive success of range-expanding species. To do this, a microcosm experiment was established where *Daucus carota* (a dryland native plant species) and *Calamagrostis epigejos* (a dryland range-expanding species) grew with soil inoculum previously conditioned by different microplastic shapes and polymer types. We expected feedback responses depending on the microplastic type with which the soil was conditioned and an increase in the performance of the range-expanding species due to the legacy effect of microplastics on soil.

## Material and methods

### Soil conditioning phase

In a previous experiment (Lozano et al. 2021b), sandy loam soil was conditioned with 12 different microplastic types. That is, soil was mixed with microplastics that had different shapes (fibers, films, foams, or fragments) and that were made from different polymer types (polyester made of at least 80% of polyethylene terephthalate (PES), polyamide (PA), polypropylene (PP), low density polyethylene (LDPE), called polyethylene from now on, polyethylene terephthalate (PET), polyurethane (PU), polystyrene (PS), and polycarbonate (PC)), at a concentration of 0.4% w/w (0.4 g of microplastic for each 100 g of dry soil). For each microplastic type, 7 replicates were established where a single seedling of *Daucus carota* grew in each pot during 4 weeks. At harvest, soil free of roots was air-dried and sampled for using in this experiment (the feedback phase). See additional details of the conditioning phase in Lozano et al. (2021b).

### Plant species selection

For the feedback phase, we selected *Daucus carota* and *Calamagrostis epigejos* as phytometers, because these species exhibit clear responses to the addition of microplastics in the soil (Lozano and Rillig, 2020; Lozano et al., 2021b). *Daucus carota* is a native biennial herbaceous plant typical of grassland ecosystems (Federal Agency for Nature Conservation, 2019) and constitutes a “conspecific” feedback, since it grows in the soil that it had conditioned; while *Calamagrostis epigejos* is a native species of range-expanding character (Těšitel et al., 2017), which appears to perform better (higher biomass) than other natives within a grassland community when microplastics are added into the soil (Lozano and Rillig, 2020) and constitutes, for this experiment, a “heterospecific” feedback, since it grows in the soil conditioned by *Daucus carota*. Seeds of these plant species were obtained from commercial suppliers in the region (Rieger-Hofmann GmbH, Blaufelden, and Jelitto Staudensamen GmbH, Schwarmstedt, Germany, respectively), surface sterilized with 4 % sodium hypochlorite for 5 min and 75 % ethanol for 2 min, thoroughly rinsed with sterile water and germinated on sterile sand. Then, seedlings of similar size were used in this experiment.

### Feedback phase experiment

In November 2019, we collected and sieved (4 mm mesh size) sandy loamy soil from Dedelow, Brandenburg, Germany (53° 37’ N, 13° 77’) where our plant species naturally grow in a diverse dry grassland. We used this soil as a substrate in microcosms (pot of 6 cm diameter, 25 cm height, 500 ml, 400 g of capacity). Then, ninety-eight seedlings of each of the two plant species were transplanted as single individuals into each microcosm, which was inoculated with soil from an independent replicate from the conditioning phase. Our experimental design included 2 plant species x 12 different soil inocula (from each of the 12 microplastic types: 4 shapes x 3 polymer types) x 7 replicates = 168 pots. Fourteen additional control pots were established per plant species. In order to prepare the soil inoculum and following Rodríguez-Echeverría et al. (2013) and Lozano et al. (2017), we took 75 g of soil from each replicate of the conditioning phase and stirred for five minutes in 150 ml of distilled, autoclaved water in a 1:2 (v:v) ratio. Then, the soil mixture was passed through a 0.5 mm sieve to remove soil particles, allowing fungal spores, hyphae, soil bacteria and microfauna to pass through (Van de Voorde et al., 2012). For the control replicates, the same procedure was followed but using substrate soil (without being conditioned with microplastics). This inoculum preparation procedure reduced any relative potential differential input of nutrients or microplastic residues with inoculation (Rodríguez-Echeverría et al., 2013). Plants in the feedback phase grew for 6 weeks. All microcosms were watered twice per week with 70 ml of water to keep water holding capacity ∼ 60 %. Plants were grown in a glasshouse chamber with a daylight period set at 12 h, 50 klx, and a temperature regime at 22/18 ºC day/night with relative humidity of ∼40 %. None of the plants died during the experiment. Microcosms were randomly distributed in the chamber and their position shifted twice to homogenize environmental conditions during the experiment.

### Measurements

At the end of the experiment, roots were carefully removed from the soil and gently washed in order to measure morphological traits in fine roots (i.e., < 2 mm in diameter which included mostly first to third order roots). We measured length, surface area, volume and root average diameter on a fresh sample using the WinRhizoTM scanner-based system (v.2007; Regent Instruments Inc., Quebec, Canada). We calculated different root morphological traits: specific root surface area (SRSA; cm^2^ mg^-1^), specific root length (SRL; cm mg^-1^), root average diameter (RAD; mm) and root tissue density (RTD; root dry weight per volume mg cm^-3^). Shoot and root mass were measured after drying samples at 70 ºC for 48 h.

### Statistical analyses

In order to test whether microplastics in soil have feedback effects on plant performance, we evaluated the effects of soil conditioned by microplastics on shoot and root masses and on root traits, through linear models and multiple comparisons (“multcomp” R package). Therefore, inoculum from soil conditioned with microplastics having different shapes and polymer types (microplastic type) were considered as fixed factors. Residuals were checked to follow assumptions of normality and homogeneity and when necessary, we implemented the function “varIdent” to account for heterogeneity. After that, we implemented the function “glht” and the “Dunnett” test from the “multcomp” R package (Hothorn et al., 2008; Bretz et al., 2011), in order to compare the effect of the soil inoculum conditioned by microplastics with the control (soil inoculum without being conditioned by microplastics). In addition, effect sizes were estimated to show the variability in the response of our variables, by comparing each inoculum conditioned by microplastics with the control pots, using a bootstrap-coupled estimation “dabestr” R package (Ho et al., 2019). Positive effects indicated that the shoot and root mass were greater with the soil inoculum conditioned by microplastics than with soil inoculum not conditioned by microplastics (positive feedback). Negative effects indicate the opposite (negative feedback), while neutral effects indicate a similar response among treatments (neutral feedback). Root trait responses were analyzed in a similar way. Positive numbers indicate a higher value of the trait with soil inoculum conditioned by microplastics than with soil inoculum not conditioned by MPs while negative numbers indicate the opposite. All data were analyzed for each plant species separately using R v.3.5.3 (R Core Team, 2019). Results shown throughout the text and figures are mean values ± 1 SE.

## Results

Microplastics in soil had a legacy effect on plant species, which depended on the microplastic shape and polymer type with which the soil was previously conditioned, and on the plant trait studied.

### Microplastic feedback effects on the native Daucus carota: Changes in biomass and root traits are evident

We found that shoot mass increased by ∼20% with soil inoculum conditioned with films, ∼17 % with foams, and ∼17 % with fragments, in comparison to the control inoculum not conditioned by microplastics (Fig. 1a, Table S1). Regarding polymer type, shoot mass increased by ∼ 35 % and ∼36 % with soil inoculum conditioned with PS foams and PET fragments, respectively (Fig. 1a, Table S2). By contrast, root mass decreased by ∼22% with soil conditioned by fibers. Regarding polymer type, it decreased by 25% and 28% with soil inoculum conditioned by PES and PA fibers (Fig.1b, Table S1, S2).

**Figure 1.**
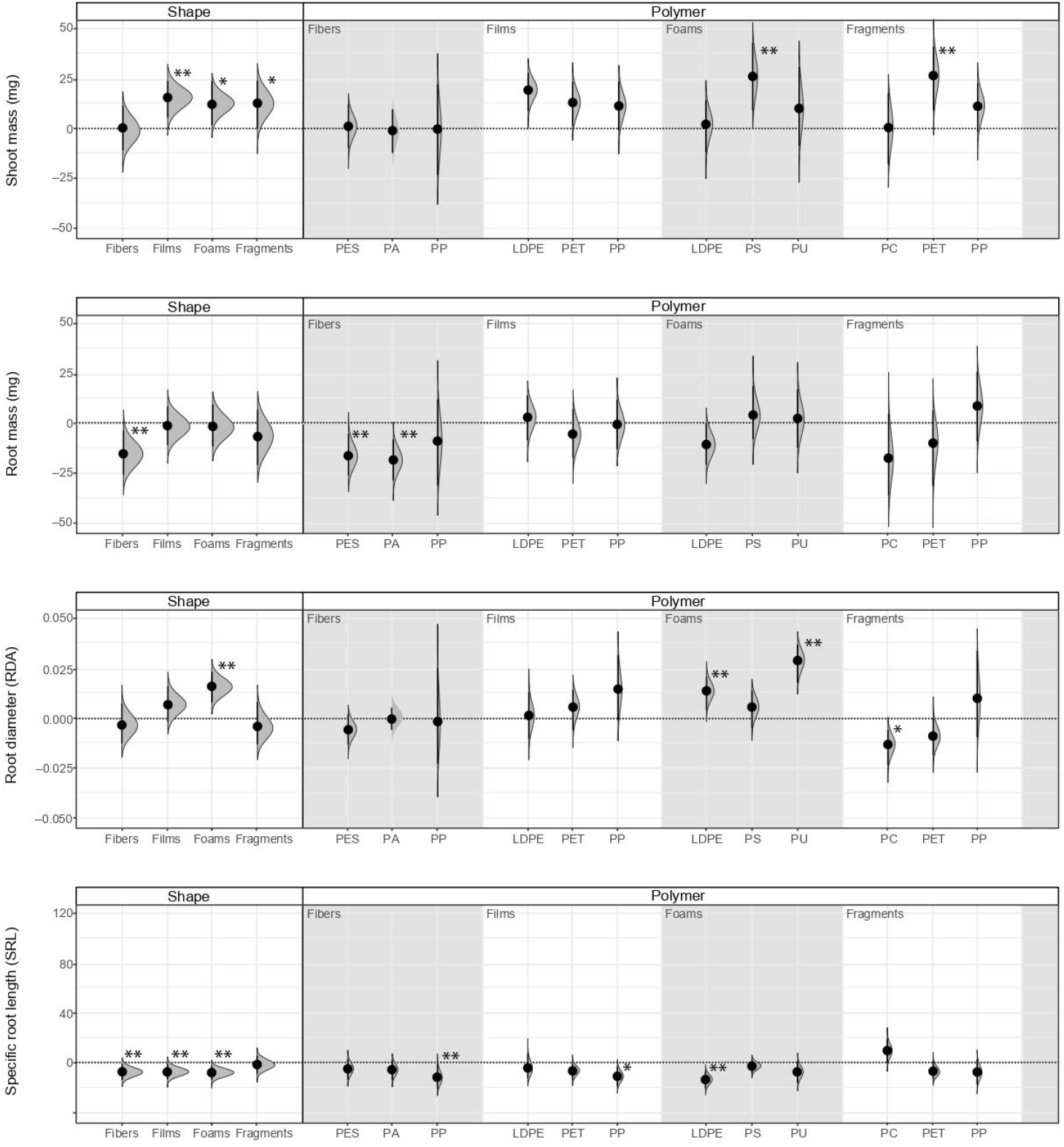
Legacy effect of microplastic shape and polymer type on shoot mass, root mass, root diameter (RDA), and specific root length (SRL) of *Daucus carota*. Effect sizes and their variance are displayed as means and 95% confidence intervals. Horizontal dotted line indicates the mean value of the control (soil conditioned without microplastics). Polymers: PES (polyester), PA (polyamide), PP (polypropylene), LDPE (low-density polyethylene), PET (polyethylene terephthalate), PS (polystyrene), PU (polyurethane), and PC (polycarbonate). Strong and moderate evidence was established at 0.05 (**) and 0.1 (*), respectively (supplementary tables S1, S2). n=7 for soil conditioned with microplastics, n=14 for control samples.

*Daucus* root morphological traits were also influenced by the legacy effect of microplastics in soil. That is, RDA increased by ∼9 % with soil inoculum conditioned with foams. Of these, it increased by ∼8% and ∼16% with soil inoculum conditioned by LDPE and PU foams, respectively. Likewise, it decreased by ∼7% with soil inoculum conditioned by PC fragments (Fig 1c, Table S1, S2). By contrast, SRL decreased with most of the microplastic shapes. SRL decreased by ∼22%, ∼21%, and ∼23% with soil inoculum conditioned by fibers, films, and foams, respectively. Of these, SRL decreased by ∼34%, ∼21%, and ∼19% with soil inoculum conditioned with PP fibers, PP films, and LDPE foams, respectively (Fig 1d, Table S1, S2).

Other root morphological traits were also affected by the legacy of microplastics in soil. RTD increased by ∼52% with soil inoculum conditioned with fibers, ∼23% with films, and ∼21% with fragments (Fig. S1a, Table S1). Regarding polymer type, RTD increased by ∼91%, ∼31%, ∼57 %, and ∼39% with soil inoculum conditioned with PP fibers, PP films, LDPE foams, and PET fragments, respectively. Similar to SRL, SRSA decreased by ∼24%, ∼19%, and ∼17% with soil inoculum conditioned by fibers, films, and foams (Fig S1b, Table S1). Of these, SRSA decreased by ∼69%, 38% and ∼54%, and 30% with soil inoculum conditioned with PP fibers, PP films, LDPE foams, and PET fragments, respectively.

### Microplastic feedback effects on the native range-expanding Calamagrostis epigejos. Effects on shoot mass are practically negligible

We did not find evidence that shoot mass of *Calamagrostis* was affected by soil inoculum as a function of having been conditioned by microplastics of different shapes. Nonetheless, regarding polymer type, shoot mass increased by ∼32% with soil inoculum conditioned with PET films (Fig. 2a, Table S1, S2). Root mass increased by ∼ 21% with soil inoculum conditioned by foams, in comparison to the control inoculum not conditioned by microplastics (Fig 2b, Table S1). Regarding polymer type, it increased by ∼33% with soil inoculum conditioned with PS foams while it decreased by ∼40 % and ∼29 % with soil inoculum conditioned by PES and PA fibers (Fig. 2b, Table S2).

**Figure 2.**
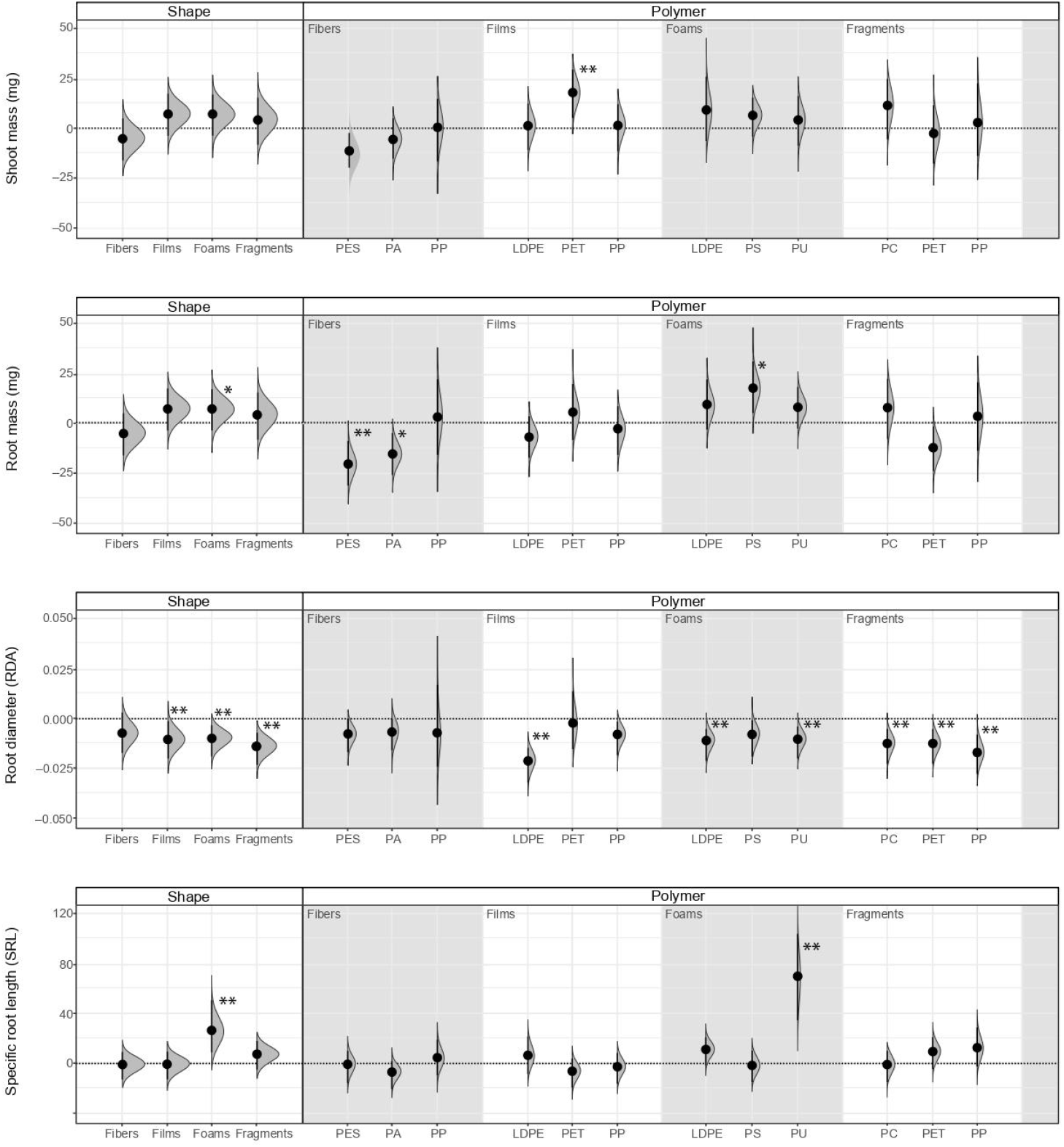
Legacy effect of microplastic shape and polymer type on shoot mass, root mass, root diameter (RDA) and specific root length (SRL) of *Calamagrostis epigejos*. Effect sizes and their variance are displayed as means and 95% confidence intervals. Horizontal dotted line indicates the mean value of the control (soil conditioned without microplastics). Polymers: PES (polyester), PA (polyamide), PP (polypropylene), LDPE (low-density polyethylene), PET (polyethylene terephthalate), PS (polystyrene), PU (polyurethane), and PC (polycarbonate). Strong and moderate evidence was established at 0.05 (**) and 0.1 (*), respectively (supplementary tables S1, S2). n=7 for soil conditioned with microplastics, n=14 for control samples.

With respect to root morphological traits, we found that RDA decreased by ∼8% with soil inoculum conditioned by films, ∼ 7% by foams and 11% by fragments (Fig. 2c, Table S1). Regarding polymer type, RDA decreased by ∼17%, ∼9 % and ∼9 % with soil inoculum conditioned by LDPE films, LDPE, and PU foams, respectively; and by ∼10%, ∼10%, and ∼14% with soil inoculum conditioned by PC, PET, and PP fragments, respectively (Fig 2c, Table S2). By contrast, SRL increased by ∼50% with soil inoculum conditioned with foams, in comparison to the control inoculum not conditioned by microplastics. Of these, SRL increased by ∼132% with soil inoculum conditioned with PU foams (Fig 2d, Tables S1, S2).

Likewise, RTD increased by ∼15 % and ∼19% with soil inoculum conditioned by fibers and films, respectively, in comparison to the control inoculum not conditioned by microplastics. Regarding polymer type, RTD increased by ∼22 % with soil inoculum conditioned with PA fibers, while by contrast, it decreased by ∼37% with soil inoculum conditioned by PU foams (Fig. S2a, Tables S1, S2). On the other hand, SRSA increased by ∼44% with soil inoculum conditioned by foams. Of these, SRSA increased by ∼123% with soil inoculum conditioned by PU foams (Fig. S2b, Tables S1, S2).

## DISCUSSION

Our results showed that microplastics in the soil had a legacy effect on soil biota with consequences for plant biomass and root morphological traits depending on plant species identity. We found that microplastics can cause a positive or negative feedback depending on the microplastic shape and polymer type that conditioned the soil.

### Microplastic films led to a positive feedback on shoot mass of *Daucus carota*

Microplastic films led to a positive feedback on shoot mass (higher mass in soil conditioned by MPs than in the control), which may be linked with microplastic films increasing soil enzymatic activities such as urease or catalase (Huang et al., 2019) as well as the abundance of nitrogen fixers bacteria (Fei et al., 2020). The priming effect of the carbon in microplastics (the addition of carbon to the soil due to MPs) may positively affect the mineralization of native soil organic C (Rillig et al., 2021), helping to explain that positive feedback. Likewise, microplastic films may lead to soil water depletion as they increase the rate of soil evaporation (Wan et al., 2019), which may alter harmful soil microbial communities in terms of diversity, abundance and activity. Indeed, previous research has found that fungal pathogens associated with *Daucus carota* decrease in abundance when soil water is reduced (Lozano et al., 2021c). A similar relationship between pathogen abundance and water reduction has been observed (Buscardo et al., 2021). Thus, microplastics films may have reduced pathogen abundance in soil or other harmful soil biota *via* effects on soil water, which, added to the promotion of mutualists, microbial activity and carbon mineralization, may help explain the positive feedback on shoot mass of *Daucus carota* caused by the legacy of microplastic films.

### Microplastic foams and fragments led to a positive feedback on shoot mass of *Daucus carota*

The positive feedback on shoot mass caused by microplastic foams and fragments may be linked with their positive effects on soil aeration (Lozano et al., 2021b) and as consequence on soil microbial activity and soil biota (Bronick and Lal, 2005). Previous research has observed that soil aeration induced by microplastics increases enzymatic activities, bacterial diversity, and the relative abundance of beneficial soil bacteria associated to nitrification and nitrogen fixation (Qian et al., 2022). Likewise, it has been shown that microplastic foams increase enzymatic activities such as cellobiosidase, β-D-glucosidase, and N-acetylβ-glucosaminidase (Zhao et al., 2021). Thus, microplastic foams and fragments may promote beneficial soil biota *via* effects on soil aeration, which in the end may help explain the positive feedback on shoot mass of *Daucus carota*.

The polymer type of which microplastic foams and fragments were made also played a role in plant-soil feedback. We observed that PS foams and PET fragments were the polymer types that led to a positive feedback on shoot mass. In that sense, previous results showed that PS increases enzyme activity such as for β-glucosidase and cellobiosidase (Awet et al., 2018) but that PS can also negatively affect other enzymes, as it is made of monomers that can be hazardous for the environment (Lithner et al., 2011).

### Microplastic fibers led to a negative feedback on root mass of both plant species

By contrast, we found that microplastic fibers had a negative feedback on root mass. Microplastic fibers may help hold water for longer increasing soil water content (De Souza Machado et al., 2019, Lozano et al., 2021b), and soil water status that appears to increase the abundance of fungal pathogens associated with *Daucus carota* (Lozano et al., 2021c). Thus, as a consequence of the legacy effect of microplastic fibers, plants might have had a decreased root mass as a consequence of pathogenic infection.

### Microplastics and their legacy effect on root morphological traits

Our results showed that the legacy effect of most microplastic shapes caused a decrease in root finesses of *Daucus carota*. From the root economic spectrum perspective, SRL and SRSA negatively correlate with root diameter (Reich, 2014, Wright et al., 2004), a situation that was most evident with microplastic foams. As mentioned, MPs foams may promote the abundance of mutualistic soil biota, a microbial group which is highly linked with root diameter (Lozano et al., 2021c). In that sense, we found that *Daucus carota* developed thicker roots with low finesses, perhaps in order to promote mycorrhizal fungi associations (Brundrett, 2002; Kong et al., 2017; Weemstra et al., 2016, Lozano et al., 2020), which support a faster nutrient acquisition (Wahl & Ryser, 2000; Withington, Reich, Oleksyn, & Eissenstat, 2006), and thus help explain the positive feedback in terms of shoot mass for *Daucus carota*. However, the legacy effect of microplastics on *Calamagrostis epigejos* was different than that on *Daucus carota*. We found that overall, microplastics’ legacy decreased root diameter of *Calamagrostis epigejos*, probably in order to diminish pathogenic infection as root diameter is strongly linked to pathogenic fungi (Lozano et al., 2021c).

### Microplastic legacy in soil affects plant species depending on their identity

We obtained strong evidence that microplastics in soil had a legacy effect on shoot mass of *Daucus carota* while the effect was not clear on shoot mass of *Calamagrosis epigejos*. Likewise, we found that root morphological traits are strongly affected by the microplastics legacy on soil and that depending on the plant species a different root trait is affected. The legacy effect of microplastics on plants are first experienced by the roots, as those are in direct contact with the soil. This is true irrespective of plant species. Then, the effect may be transferred to plant biomass, depending on the plant species. For example, unlike *Daucus carota*, which was affected in its root traits and plant biomass by microplastics legacy, *Calamagrostis epigejos* was affected in its root traits but not in its plant biomass, which shows that microplastics effects on plants are species-specific, as is the case with other global change factors (Lozano et al., 2020). This showed that, at least in the short term, microplastics legacy on soil does not contribute to the competitive success of this range-expanding species.

Our results showed that microplastics have a legacy effect on plant biomass and root morphological traits which can be positive or negative as a function of microplastic shape and polymer type. Certainly, the positive feedback does not mean a desirable effect but simply an increase in plant biomass and alterations in root morphological traits. Indeed, these positive effects still represent deviations from the natural state (Rillig et al. 2021). Our study provides novel insights into the effects of microplastics on terrestrial systems highlighting their key role in plant-soil feedbacks.

## ACKNOWLEDGEMENTS

The work was funded by the German Federal Ministry of Education and Research (BMBF) within the collaborative Project “Bridging in Biodiversity Science (BIBS)” (funding number 01LC1501A). M.C.R. additionally acknowledges support from the EU grants MINAGRIS and PAPILLONS. We thank Matti Skarupke, Andrea C. Gundry and Emily Magkourilou for their help in the establishment of the experiment and data collection. We also thank Sthefania Quintero-Pineda for her help in the figures design.

## AUTHOR CONTRIBUTIONS

YML conceived the ideas and designed the methodology with input from MCR; YML established and maintained the experiment in the greenhouse, analyzed the data, and wrote the first draft. MCR contributed to the draft with his comments and editions. Both authors gave final approval for publication. We have no conflict of interest to declare.

## DATA AVAILABILITY

Data that support the findings of this study are available from the corresponding author upon request.

## Supplementary material

**Table S1.**
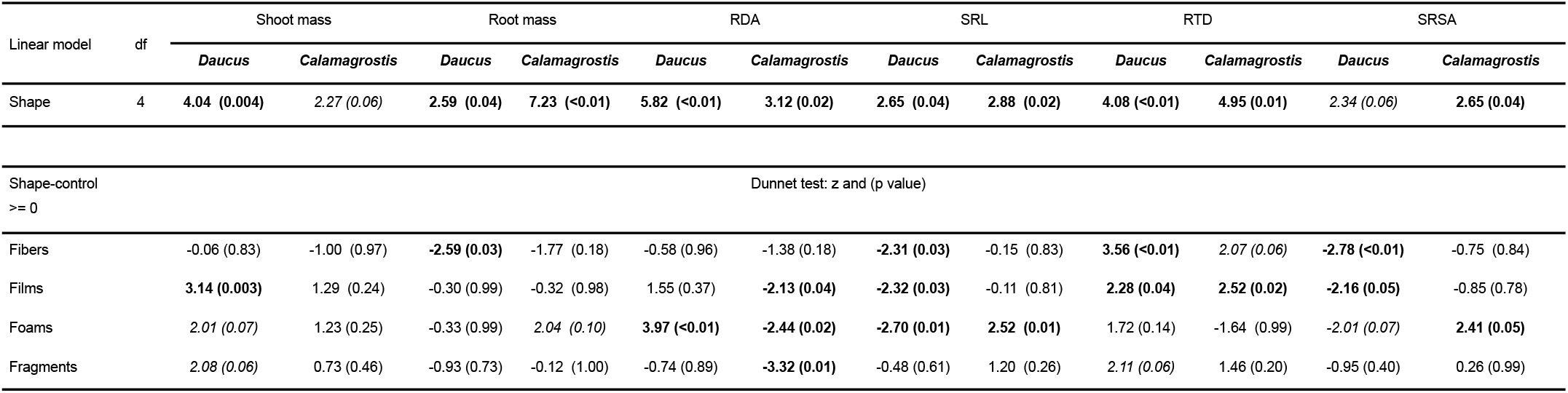
Legacy effect of microplastic shape on plant mass and root traits. Results of linear model (F and p value) and multiple comparisons by using the Dunnett test: z and (p value). Values in bold signify a strong effect (p<0.05) and, in *italic* a moderate effect (p<0.1) of the treatment on the dependent variable.

**Table S2.**
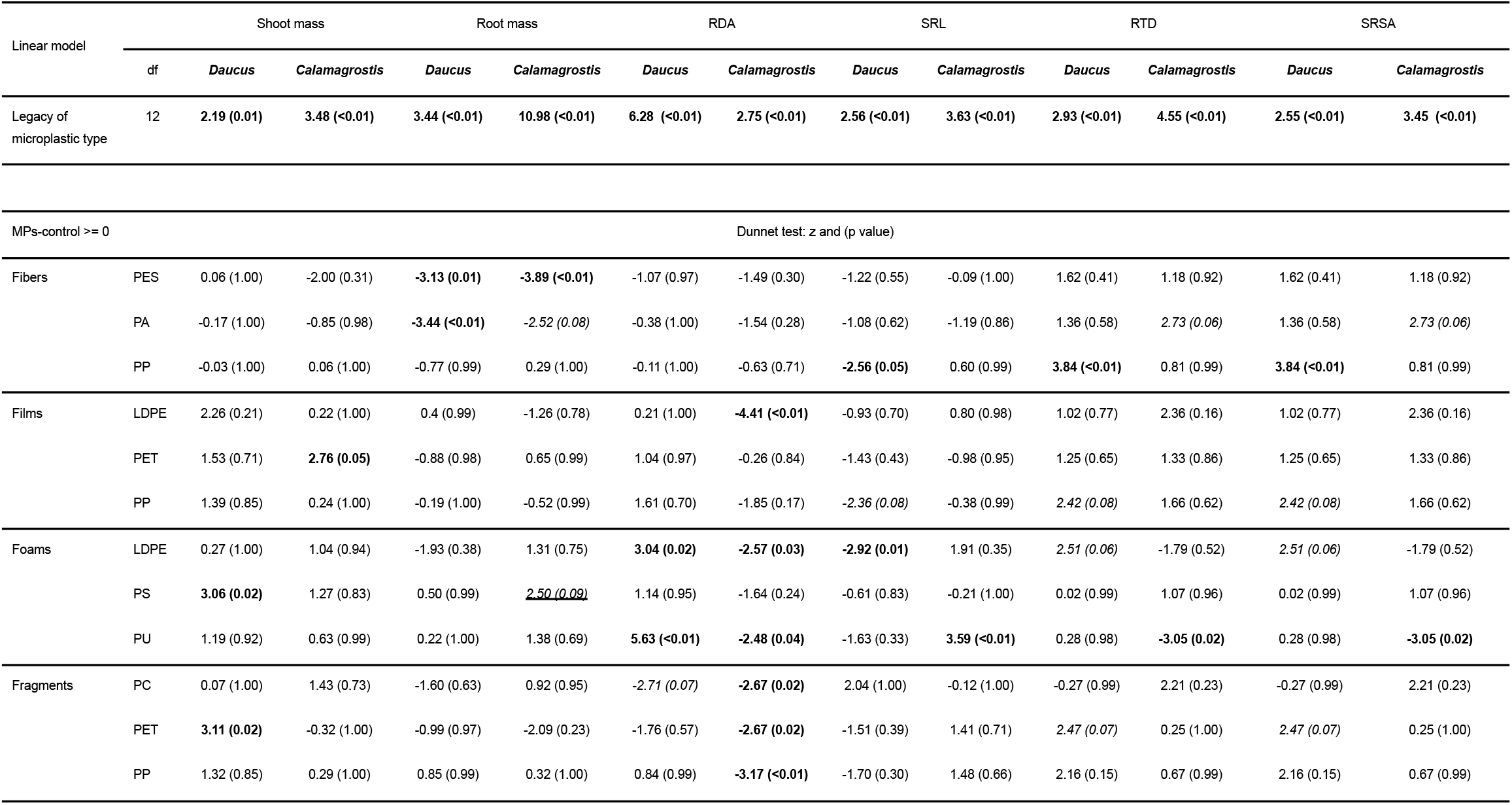
Legacy effect of microplastic type effect on plant mass and root traits. (F and p value) and multiple comparisons by using the Dunnett test: z and (p value). Polypropylene (PP); Polyester (PES); Polyamide (PA); Polyethylene (PE); Polyethylene terephthalate (PET); Polyurethane (PU); Polystyrene (PS); Polycarbonate (PC). Values in bold evidence a strong effect (p<0.05), and in *italic* a moderate effect (p<0.1) of the treatment on the dependent variable.

**Figure S1.**
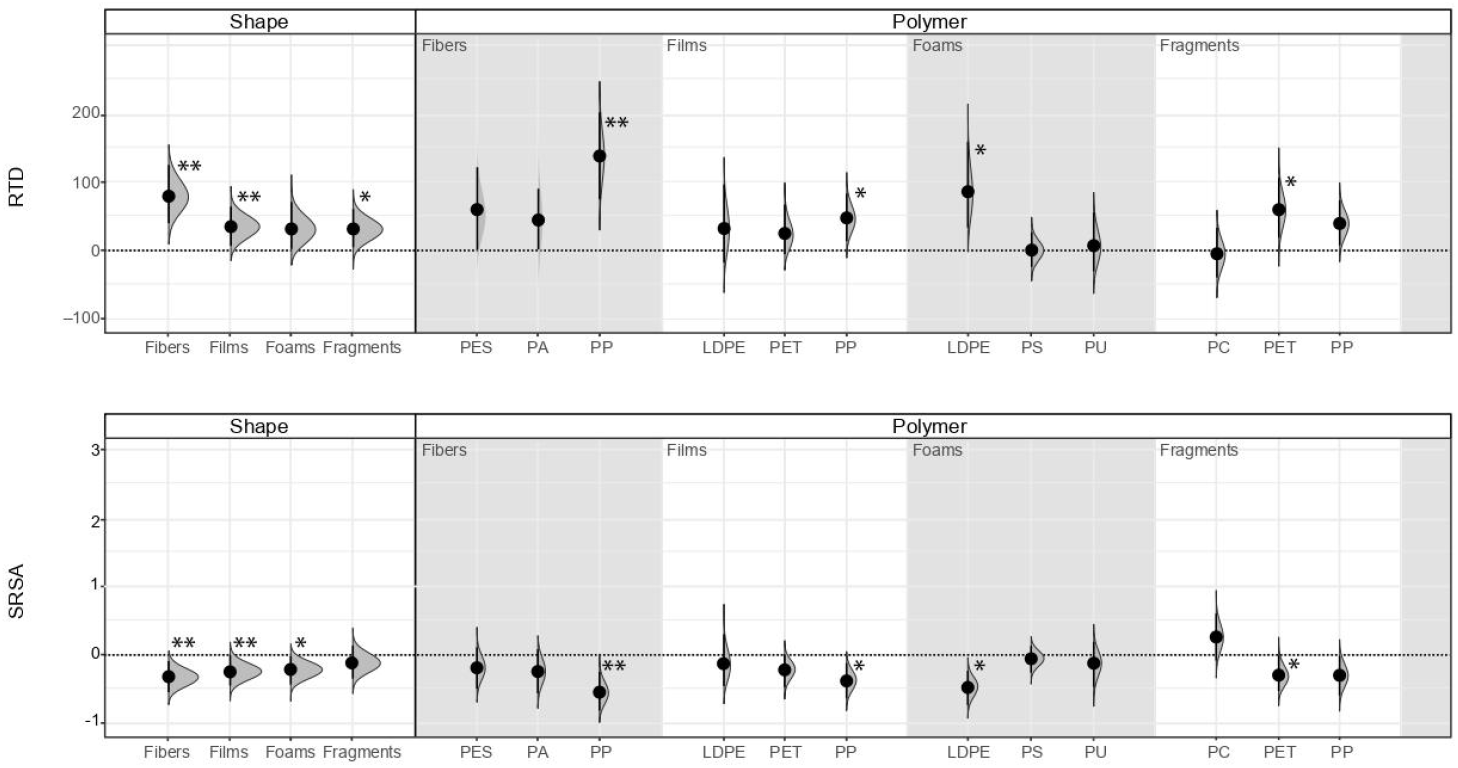
Legacy effect of microplastic shape and polymer type on root tissue density (RTD) and specific root surface area (SRSA) of *Daucus carota*. Effect sizes and their variance are displayed as means and 95% confidence intervals. Horizontal dotted line indicates the mean value of the control (soil conditioned without microplastics). Polymers: PES (polyester), PA (polyamide), PP (polypropylene), LDPE (low-density polyethylene), PET (polyethylene terephthalate), PS (polystyrene), PU (polyurethane), and PC (polycarbonate). Strong and moderate evidence was established at 0.05 (**) and 0.1 (*), respectively (supplementary tables S1, S2). n=7 for soil conditioned with microplastics, n=14 for control samples.

**Figure S2.**
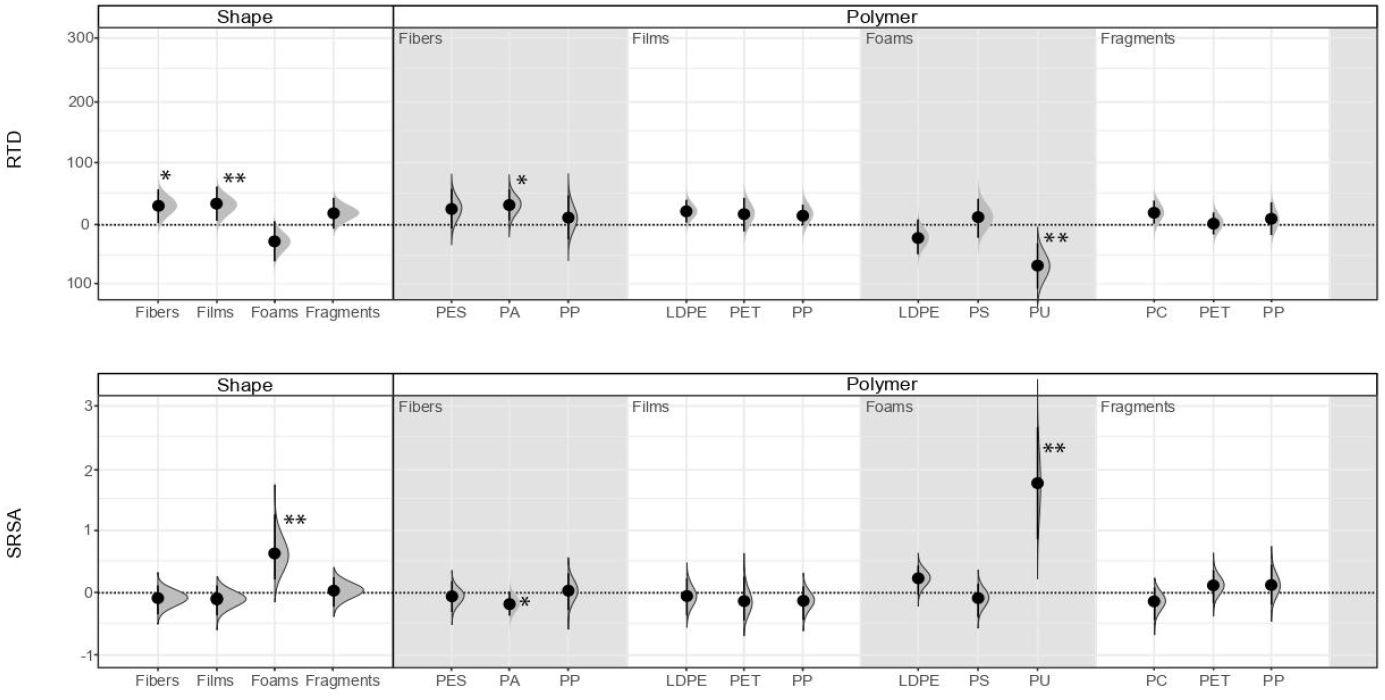
Legacy effect of microplastic shape and polymer type on root tissue density (RTD) and specific root surface area (SRSA) of *Calamagrostis epigejos*. Effect sizes and their variance are displayed as means and 95% confidence intervals. Horizontal dotted line indicates the mean value of the control (soil conditioned without microplastics). Polymers: PES (polyester), PA (polyamide), PP (polypropylene), LDPE (low-density polyethylene), PET (polyethylene terephthalate), PS (polystyrene), PU (polyurethane), and PC (polycarbonate). Strong and moderate evidence was established at 0.05 (**) and 0.1 (*), respectively (supplementary tables S1, S2). n=7 for soil conditioned with microplastics, n=14 for control samples.

